# Identifying critical transcriptional targets of the MYC oncogene using a novel competitive precision genome editing (CGE) assay

**DOI:** 10.1101/2021.09.17.460746

**Authors:** Päivi Pihlajamaa, Otto Kauko, Biswajyoti Sahu, Teemu Kivioja, Jussi Taipale

**Author notes:** Corresponding author: Jussi Taipale.

## Abstract

MYC is an oncogenic transcription factor that controls major pathways promoting cell growth and proliferation. MYC has been implicated in the regulation of large number of genes, but the exact target genes responsible for its proliferative effects are still not known. Here, we use a novel competitive genome editing (CGE) assay for studying the functional consequence of precise mutations of MYC binding sites on cell proliferation. The CGE method is based on precision genome editing, where a CRISPR/Cas9-induced DNA break is repaired using a template that either reconstitutes the original feature or introduces an altered sequence. Both types of repair templates harbor sequence tags that allow direct comparison between cells that carry original and mutant features and generate a large number of replicate cultures. The CGE method overcomes the limitations of CRISPR/Cas9-technology in analyzing the effect of genotype on phenotype, namely the difficulty of cutting DNA exactly at the intended site, and the decreased cell proliferation caused by the DNA cuts themselves. Importantly, it provides a powerful method for studying subtle effects elicited by mutation of individual transcription factor binding sites. We show here that E-box mutations at several MYC target gene promoters resulted in reduced cellular fitness, demonstrating a direct correlation between MYC-regulated cellular processes and MYC binding and identifying important transcriptional targets responsible for its functions.

## Introduction

MYC is a basic helix-loop-helix (bHLH) transcription factor (TF) that forms a heterodimer with another bHLH protein MAX and regulates a large set of target genes by binding to a CACGTG sequence (E-box) at their promoters^1,2,3^. MYC is indispensable for embryonic development^4^ but in normal cells its expression is tightly controlled. The importance of tight regulation of MYC activity is highlighted by the fact that it is one of the most frequently deregulated oncogenes across multiple human cancer types^5^. MYC regulates major pathways promoting cell growth and proliferation, such as ribosome biogenesis and nucleotide biosynthesis^6^. However, dissecting primary transcriptional targets of MYC from secondary effects among the large number of potential targets has been challenging. It has been proposed that MYC, instead of being a regulator of a particular transcriptional programs, is a universal amplifier of gene expression that increases transcriptional output at all active promoters^7,8^. Conversely, it has been shown that MYC can selectively regulate specific sets of genes, including those involved in metabolism and assembly of the ribosome^9,10^. Nevertheless, despite its well-known phenotypic effects on cellular growth and proliferation, the precise MYC target genes accounting for its oncogenic activity are still elusive.

The most effective way to dissect the gene regulatory network downstream of MYC would be to individually assess the role of each target gene by mutating the MYC binding sites at its promoter. The current genome editing tools such as bacterial clustered regularly interspaced short palindromic repeats (CRISPR)–CRISPR-associated protein 9 (Cas9) have proven to be robust and efficient tools for many sequence manipulations. They have been extensively used for mutating specific genomic loci in single-gene studies^11^ as well as genome-wide screens^12,13,14^. However, resolution of the CRISRP/Cas9-editing is limited by the suitable protospacer adjacent motif (PAM) sequences found in the close proximity of the region of interest. Homology-directed recombination (HDR)-mediated precision editing can be used to introduce genetic alterations exactly at the intended loci, but this method suffers from strong DNA damage response, low efficiency, and the incompatibility with pooled CRISPR-screening approaches. Because of the low efficiency of precision genome editing, pooled screens commonly use lentiviral introduction of libraries of guide-RNAs to cell lines that express either Cas9 nuclease alone that generates a series of insertion and deletion alleles, or nuclease dead Cas9 fused to a transcriptional repressor (CRISPRi) or activator (CRISPRa) elements^15,16,17^. These methods do not have single base or single allele resolution, and their precision is limited because they use an indirect measure, inferring the perturbation from the presence of a guide-sequence integrated into the cells at a (pseudo)random genomic position.

Furthermore, interpreting the functional consequence of targeted Cas9-induced mutations is confounded by the DNA damage introduced by Cas9, and from off-target effects of the Cas9 nuclease^18^. In particular, double-strand breaks (DSB) at on- or off-target loci cause DNA damage and genomic instability resulting in paused cell cycle or apoptosis^19,20,21^. These problems are particularly acute in analysis of small intergenic features, such as transcription factor binding sites. This is because sgRNAs cannot be selected from a large number of possible sequences predicted to have the same effect, and the flanking sequences of most TF binding sites are generally less complex than sequences found in coding regions. Here we describe a novel competitive precision genome editing (CGE) approach utilizing CRISPR/Cas9 genome editing at precise loci to accurately analyse the effect of mutations on cellular properties and molecular functions, such as fitness, transcription factor (TF) binding, or mRNA expression. Importantly, the novel experimental design in the CGE approach mitigates the confounding factors associated to CRISPR experiments, such as the hampering effect of double-strand DNA break itself on cell proliferation, enabling dissection of the effect of regulation individual MYC target genes on cellular fitness.

## Results

### A novel CGE approach enables lineage-tracing and quantitative analysis of phenotypic effects elicited by precise mutations

The novel CGE method utilizes CRISPR/Cas9 technology combined with one or several HDR templates with sequence tags enabling lineage-tracing of the targeted cell populations. One of the HDR templates (control) reconstitutes the wild-type sequence of the region of interest by harboring the original genomic sequence, whereas the other template will replace it with desired mutated sequence, such as non-functional TF binding site (**Fig. 1a**). Importantly, each individual HDR template molecule has variable sequence tag(s) flanking the sequence of interest that can be detected from the Illumina sequencing reads of the targeted locus (**Fig. 1a**). This enables precise counting of the editing events facilitated by each repair template and direct comparison of the effect of the mutation to original genomic sequence. In addition, inclusion of a large set of different sequence tags allows excluding the possibility that the tags themselves, and not the intended mutations, cause the observed phenotype.

**Figure 1.**
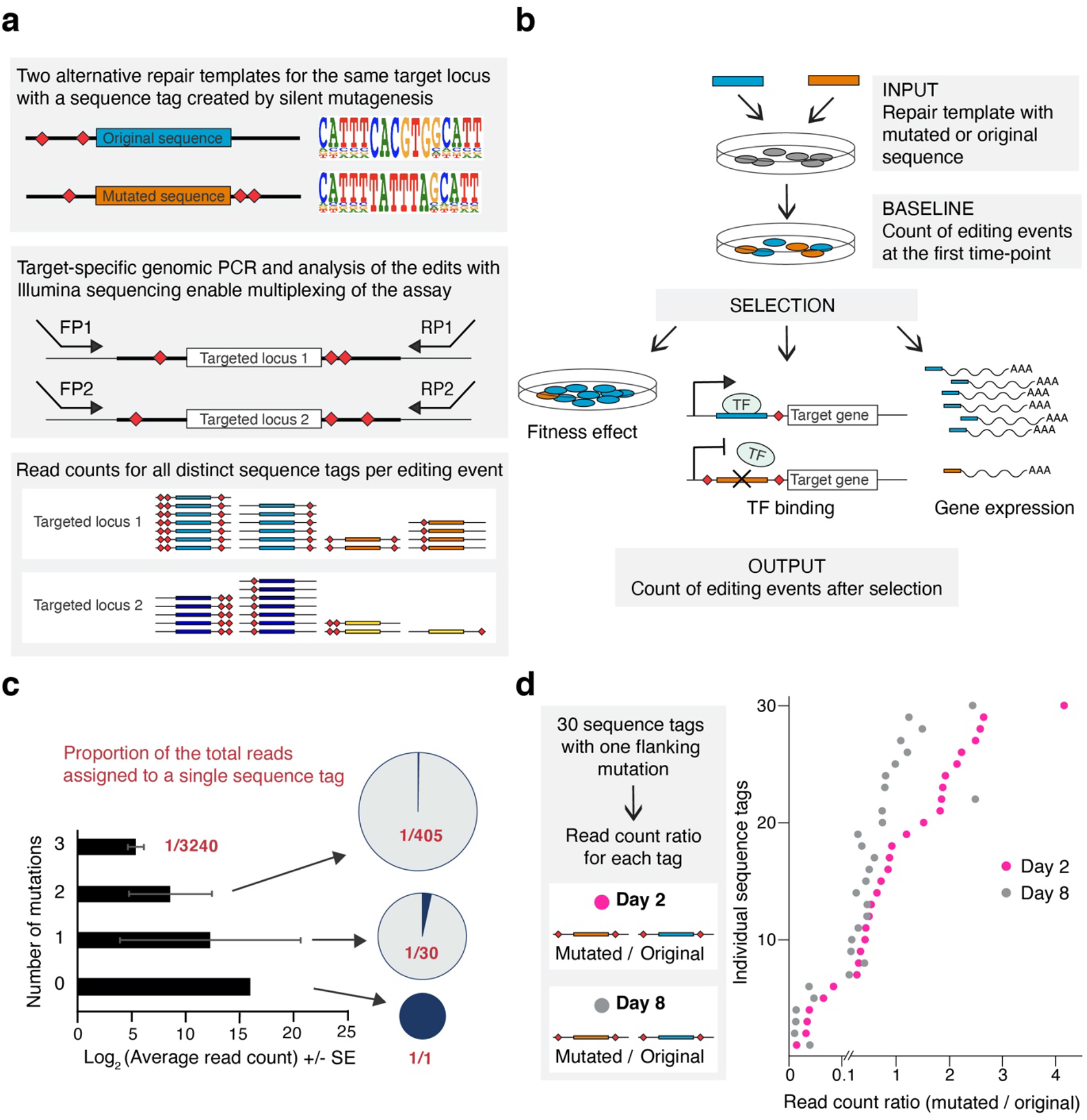
Strategy for lineage-tracing of cells with distinct genome editing events utilizing sequence tags with silent mutations. **a**, Conceptual outline of the precision editing strategy utilizing two HDR templates, one harboring original genomic sequence and another with the desired mutation. Five nucleotides on both side flanking the region of interest are mutated in the HDR oligos with the probability of 24%, a strategy that maintains most of the flanking sequence intact. Editing events at targeted locus can be identified using Illumina sequencing and cell lineages traced according to their mutation signatures. **b**, Schematic presentation of the experimental workflow utilizing a 1:1 mixture of two HDR templates for the same target. The abundance of each HDR template in the cell population can be analyzed at the baseline (first time point), and after selection pressure using different output assays, such as cellular fitness (after one week of culture), TF binding (after immunoprecipitation with specific antibody), and mRNA expression. **c**, Number of distinct sequences with zero, one, two, and three mutations in the ten flanking nucleotides when mutated with the probability of 24% and their average read counts from the Illumina sequencing reads at the targeted SHMT2 locus at baseline (from ChIP input sample). Pie charts illustrate the proportion of the reads assigned to each distinct sequence tag with different number of mutations. **d**, The effect of E-box mutation at the *RPL23* gene promoter on fitness of HAP1 cells. Precision editing results are shown separately for each cell lineage harboring a sequence tag with exactly one flanking mutation. Read count ratios for mutated vs. original sequence are shown in two time points, day 8 (grey) and day 2 (pink).

In the CGE experiment, DNA samples from cells edited with either mutant or control sequence are collected at two or more time points (early and late) and the cell lineages with particular editing event can be followed before and after subjecting the cells to selection pressure, such as competitive growth in culture, after which cellular fitness, TF binding to the target locus, and the expression levels of mRNA can be analyzed (**Fig. 1b**). Since the sequence tags are present in both repair templates, this experimental design allows precise comparison of the mutated vs. control sequence by excluding the non-edited wild-type sequences from the analysis. Sequencing reads will then be assigned to the distinct editing events based on their sequence tags, and the ratio between mutated and control sequences for each tag can be determined at both time points resulting in dozens of replicate measurements for each editing event (**Fig. 1a**). Thus, statistical power to detect differences between the time points is very high. The experiment is a single-well assay in which both the repair templates are transfected to cells within one culture well and the genomic perturbation is compared directly to control in the same cell population. This eliminates the experimental bias and variation coming from transfection/transduction and Cas9 introduced DSBs, and well-to-well variation caused by differences in culture and experimental conditions between wells.

To preserve potentially functional flanks of the sequence of interest, it is important to introduce silent or near-silent mutations. For coding regions, this can be accomplished by introducing synonymous mutations of codons and avoiding splice-junctions. Since less is known about functional elements within non-coding regions, we decided to use a diverse library that largely conserves the wild-type sequence, introducing only one or few point mutations per cell within a region wider than a typical TF binding site (~ 10 bp). In our case, each of the ten positions within the sequence tag was mutated with probability of 24%, thus keeping most of the flanking sequence intact (**Fig. 1a**) but introducing typically 2-3 mutations per repair oligo (in ~53% of the sequences; **Supplementary Fig. 1a**). This generates 30 distinct sequences whose sequence differs from the native sequence by exactly one nucleotide (**Supplementary Fig. 1b**), 405 distinct sequences with 2-nt difference to the native sequence, and 3240 distinct sequences with three mutations. In the oligo synthesis for HDR templates, the probability for any individual sequence tag with one mutation is higher than for tag with two or three mutations, which is reflected in the data with single-mutation tags having higher read counts than double and triple mutants (**Fig. 1c**), consistent with the fact that single-mutation sequence tags are present in the original mixture of synthesized oligos in more copies than double- and triple-mutants. After assigning the read counts for each editing event with mutated or native sequence at the two experimental time-points, the ratio of mutated / native sequences can be plotted for each sequence tag separately, enabling robust and accurate measurement for the effect of the mutation on cellular fitness for each cell lineage separately (**Fig. 1d**).

### CGE approach reveals protein phosphorylation sites critical for cellular fitness

To validate our novel CGE approach in functional studies, we first introduced mutations to the coding regions of genes. To this end, we mutated previously described phosphorylation sites of the cyclin-dependent kinase 1 (*CDK1*) and the growth factor receptor-binding protein 2 (*GRB2*) genes. In coding regions, sequence tags were generated by randomizing the degenerate positions of the adjacent codons in the repair template. Phosphorylation sites were abolished by alanine (A) or phenylalanine (F) substitutions of the phosphorylated serine (S), threonine (T) or tyrosine (Y) residues. To mimic phosphorylation, the same amino acids were also mutated to the acidic residues glutamate (E) or aspartate (D), which in many proteins can lead to the same effect as phosphorylation of the serine, threonine and tyrosine residues^22^. In the CGE assay, the cell lineages that carry mutations that impair cell growth should be underrepresented in the cell population after one week of culture compared to cells edited with wild-type sequence and the sequence tag, which can be analyzed from genomic DNA collected at the beginning and at the end of the experiment (**Fig. 1b**). The experiments were carried out in haploid HAP1 and near-haploid chronic myelogenous leukemia KBM-7 cell lines. HAP1 cells are a derivative of KBM-7 that grow adherently and no longer express hematopoietic markers. Most of the HAP1 cells in early passage cultures are haploid for all chromosomes, and thus particularly useful for mutational screens since only one editing event is sufficient for a full knock-out.

Previous mutagenesis screen by Blomen et al. (ref. ^23^) suggests that BCR/ABL-GRB2-RAS/MAPK signal transduction is essential for KBM-7 but not for HAP1 cells. To test this, we mutated two key residues, Y160 and Y209 in the adaptor protein GRB2 that links tyrosine kinase signaling to the RAS-mitogen-activated protein kinase (MAPK) pathway. GRB2 phosphorylation at these residues have opposing functions, phosphorylated Y209 activating and Y160 inhibiting MAPK signaling^24,25^. In agreement with this, we observed that KBM-7 but not the HAP1 cells were sensitive to GRB2 mutation (**Supplementary Fig. 2**), indicating that the precision genome editing assay can be used to identify functionally important phosphorylation events in cells.

To further validate the CGE assay we evaluated the fitness effect of CDK1 regulatory phosphorylation site mutations in human cells. CDK1 activation and onset of mitosis requires phosphorylation of T161 in the activation segment, and dephosphorylation of T14 and Y15^26^. The non-phosphorylatable double-mutant T14A/Y15F cells were almost completely lost after one week of precision editing (**Fig. 2a**). These findings are consistent with earlier work reporting that the T14A/Y15F double-mutant can be activated prematurely during the cell cycle^27^, and overexpression of this mutant in cells results in cell death due to mitotic catastrophe^28^. The effect of the phosphorylation site mutation in the CDK1 activating segment, T161A, was less prominent. Loss of phosphorylation resulted in markedly decreased cell proliferation, whereas T161E phosphomimetic mutation allowed cells to proliferate normally (**Fig. 2a**). This is consistent with the lack of requirement of regulation of the CDK activating kinase in human cells^29^. We also tested recently reported prime editing method^30^ for mutating a phosphorylation site and for introducing the sequence tag within the *CDK1* coding region. Using this approach, we observed reduced fitness of HAP1 cells as a result of Y15F mutation (**Fig. 2a**), demonstrating that also prime editing can be utilized for generating the targeted mutations and sequence tags for our precision genome editing assay.

**Figure 2.**
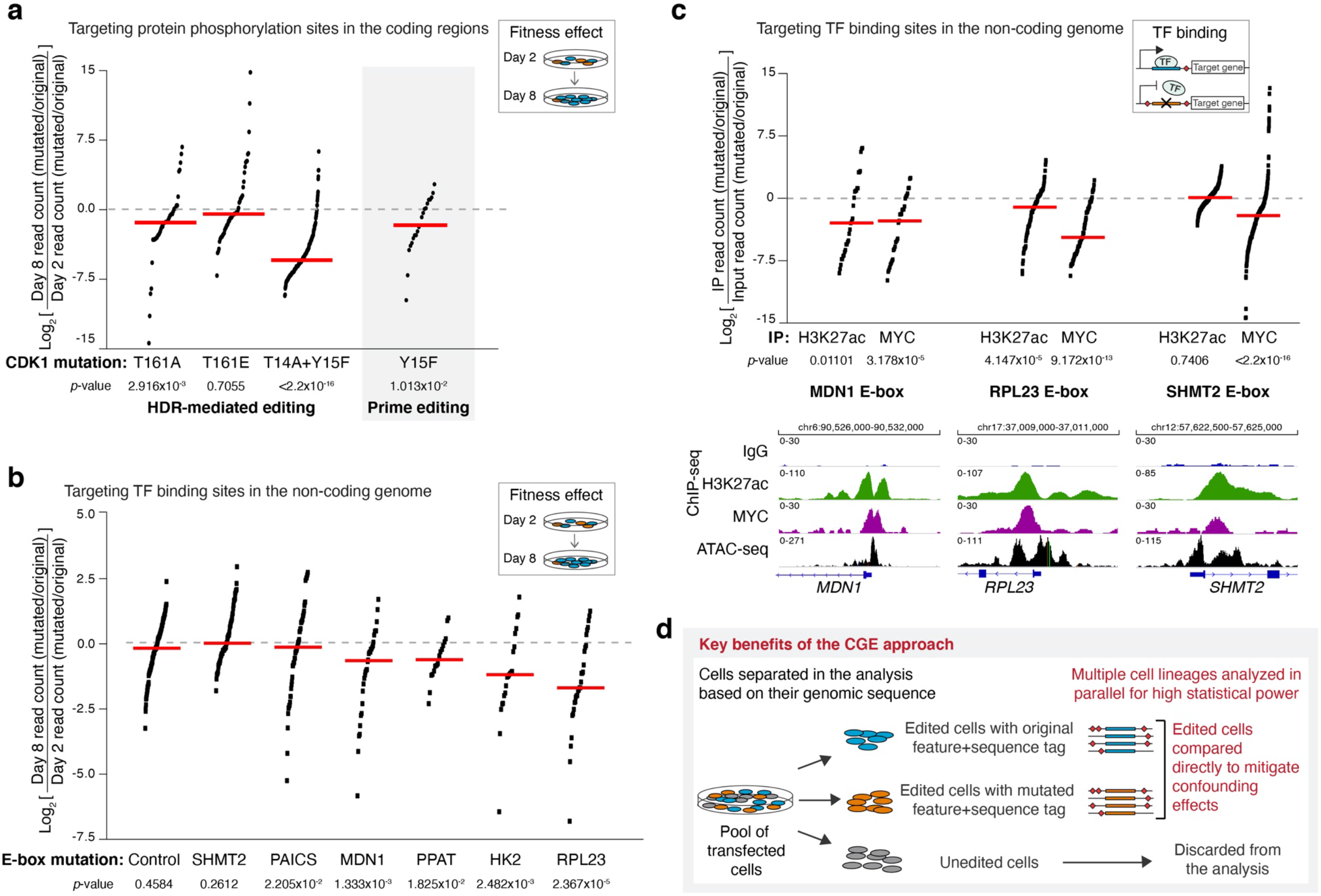
The effect of mutating TF binding sites and protein phosphorylation sites on cellular fitness determined by lineage-tracing of editing events. **a**, The effect of mutating protein phosphorylation sites in the *CDK1* gene on fitness of HAP1 cells. Precision editing results are shown for all cell lineages harboring sequence tags with read count >5 on day 2. Log_2_ values for day 8/day 2 ratios are shown for each sequence tag pair after calculating the ratio of read counts for mutated vs. original sequence at both timepoints. Note that T161E is a phosphomimetic mutation whereas T161A abolishes the phosphorylation site. Phosphorylations at T14 and Y15 inhibit CDK1 function, but the presence of non-phosphorylatable T14A/Y15F double-mutant can result in premature mitosis and mitotic catastrophe^28^. The effect of Y15F mutation on the fitness was also measured after introducing this mutation to the HAP1 cells using prime editing^30^. In panels **a-c**, red lines represent the median values, and *p*-values from Wilcoxon signed rank test are shown for each experiment (see **Supplementary Table 3** for details of the statistical parameters). **b**, The effect of mutating MYC binding motifs (E-box) at promoters of MYC target genes on fitness of HAP1 cells. E-box mutation at six MYC target gene promoters (see **Supplementary Fig. 3** for E-box locations) as well as negative control region (synonymous mutation in the MYC coding region) are shown. Precision editing results are shown for all cell lineages harboring sequence tags with exactly two flanking mutations with read count >50 on day 2. Log_2_ values for day 8/day 2 ratios are shown for each sequence tag pair after calculating the ratio of read counts for mutated vs. original sequence at both timepoints. **c**, The effect of E-box mutation on MYC occupancy and acetylation of H3K27 at promoters of MYC target genes *MDN1*, *RPL23*, and *SHMT2*. ChIP using MYC and H3K27ac antibodies followed by target-specific PCR and Illumina sequencing was performed 48 h after RNP transfection. Input from crosslinked and sonicated chromatin was used as a control. Precision editing results are shown for all cell lineages harboring sequence tags with exactly two flanking mutations with read count >50 in the input. Log_2_ values for immunoprecipitated (IP) sample/input ratios are shown for each sequence tag pair after calculating the ratio of read counts for mutated vs. original sequence in both timepoints. Genome browser snapshots show ChIP-seq and ATAC-seq tracks from wild type HAP1 cells for each of the targeted loci, demonstrating robust MYC binding to these sites in HAP1 cells. **d**, Schematic presentation highlighting the key advantages of the CGE method: high statistical power from analyzing multiple cell lineages within a single experiment, and mitigation of the confounding effects characteristic to CRISPR/Cas9-mediated genome editing, namely off-target effects, and the effect of double-strand break itself of cell proliferation, by excluding the unedited cells from the analysis.

### The CGE approach identifies precise MYC regulatory events that promote cell proliferation

After demonstrating the power of the precision editing approach in studying the functional consequence of individual protein phosphorylation sites, we utilized it for studying the gene regulatory elements within the non-coding genome. Specifically, a six-nucleotide MYC binding motif (E-box) was mutated at the promoters of MYC-target genes to study their effect on cell proliferation and fitness. If a particular E-box is essential for cell growth, the alleles containing tags and the wild-type sequence should be enriched in the cell population compared to the E-box deleted alleles after one week of culture (**Fig. 1b, d**). Although a large number of genes have been reported as MYC target genes^6^, the functional consequence for cell proliferation resulting from MYC binding to a promoter of a particular gene has not been previously shown. For the purpose of this study, putative MYC target genes were selected for editing on the basis of functional genomics studies in human colon cancer cell lines and previously published data sets in HAP1 haploid cell line using the following criteria: (1) Gene must be essential in HAP1 cells, i.e. found in both publications (refs ^23,31^), and (2) display robust MYC binding at its promoter within open chromatin on the basis of ATAC-seq signal, and clear change in expression upon MYC silencing in colon cancer cells (ref. ^32,33^; **Supplementary Fig. 3**). In addition, (3) the gene should preferably only contain one E-box within the ChIP-nexus peak (ref. ^32^; **Supplementary Fig. 3**).

The CGE experiments were carried out in HAP1 cells, and the cell lineages harboring either the original or mutated sequence were analyzed at day 2 and day 8. Importantly, targeted mutation of the E-box sequence to a non-functional TATTTA at the promoters of four MYC target genes – ribosomal protein L23 (*RPL23*), hexokinase 2 (*HK2*), phosphoribosyl pyrophosphate amidotransferase (*PPAT*), and midasin AAA ATPase 1 (*MDN1*) – resulted in reduced cell growth as measured from the read counts for lineage-tracing sequence tags with two mutations at day 8 as compared to day 2 (**Fig. 2b**). However, there were E-boxes at promoters of MYC target genes that can be mutated to non-functional sequence without affecting cell proliferation, such as serine hydroxymethyltransferase 2 (*SHMT2*) and phosphoribosylaminoimidazole carboxylase and phosphoribosylaminoimidazolesuccinocarboxamide synthase (*PAICS*; **Fig. 2b**), demonstrating the strength of this approach in dissecting the contribution of each individual transcription factor binding site to cell proliferation. Furthermore, the CGE assay can robustly measure the effect of each E-box on cellular fitness also for genes that harbor several of them within their regulatory region, as demonstrated for the *MDN1* gene. Out of the two E-boxes within the *MDN1* promoter, mutation of the E-box closer to the TSS (TSS +32) had an effect on cell proliferation, whereas the mutation of the E-box farther away (TSS-151) had no effect, despite MYC binding detected at both of these sites using ChIP-nexus (ref. ^32^; **Supplementary Fig. 4**).

Since the competitive precision genome editing assay showed clear effects on cell proliferation resulting from a mutation of a single MYC binding motif, we set to analyze the direct effects of E-box mutation on MYC binding to the promoter, and activation of the promoter as measured by an increase in the active chromatin mark histone 3 lysine 27 acetylation (H3K27ac). For this, we performed chromatin-immunoprecipitation using anti-MYC and anti-H3K27ac antibodies from the HAP1 cells after precision editing. To quantify the editing events, each targeted locus was amplified using PCR and the amplicons were Illumina sequenced. Importantly, we detected fewer antibody-enriched sequences with TATTTA mutated sequence compared to CACGTG original sequence, demonstrating less MYC binding to the mutated sequences at RPL23, MDN1, and SHMT2 E-boxes, as opposed to the input sample with equal ratios of TATTTA and CACGTG (**Fig. 2c**). We also observed decrease in H3K27 acetylation at RPL23 and MDN1 E-boxes (**Fig. 2c**), suggesting lower expression of these target genes. However, there was no changes in the level of H3K27 acetylation at the SHMT2 locus, consistent with the observation that mutation of this E-box had no effect on cell proliferation (**Fig. 2b, c**). In conclusion, we identified here several genes that are directly regulated by MYC and demonstrate that mutation of a single MYC binding motif is sufficient for reducing cellular fitness.

## Discussion

Here, we show a novel method for precise analysis of the effect of mutations on cellular phenotype by utilizing CRISPR/Cas9 precision editing combined with lineage-tracing sequence tags in HDR templates, and employ it for studying the precise effects of individual transcription factor binding sites and posttranslational modifications. Previously, next-generation sequencing based methods, such as GUIDE-seq, have been developed for assessing the off-target DNA cleavage sites^34^ and random sequence labels have been used for increasing precision and accuracy of CRISPR-screens^35^. In a recent saturation mutagenesis screen, a single-nucleotide variants (SNV) targeting BRCA1 gene were transfected to target cells along with Cas9 and sgRNA and targeted gDNA and RNA sequencing was performed to quantify SNV abundances^36^. However, our approach enables for the first time a precise assessment of the effect of particular intended mutations. Our approach of using parallel editing of the target loci with two HDR templates in a single cell culture has two key advantages over previously described genome-editing assays (**Fig. 2d**). First, silent mutations that generate sequence tags to HDR templates provide means to discard all confounding information from the next-generation sequencing output of the assay. Second, direct comparison of the mutated sequence to the reconstituted native sequence mitigates all the detrimental off-target effects, as well as enables lineage tracing of edited clones thus providing statistical power to the analysis. When measuring allele-specific phenotypes, the method also allows the use of diploid cells for analysis of phenotypes such as TF binding or RNA expression. Measuring more complex phenotypes in diploid cells is also possible, but requires either prior deletion of one allele from the targeted locus, or dilution of the two repair templates by a template that inactivates the wild-type allele in such a way that most cells carry either two inactive alleles, or one inactive allele, and one targeted allele. This will be easier when targeting coding regions, as failure of targeted repair commonly leads to inactivation of the target gene due to generation of frameshift or deletion alleles by non-homologous end-joining.

The CGE method is particularly useful for studying the effect of small sequence features such as individual transcription factor binding sites and posttranslational modifications as shown here for MYC binding motifs and phosphorylation sites in the CDK1 and GRB2 proteins, since precision editing is not dependent on finding a highly specific guide sequence precisely overlapping the feature of interest. In addition, the phenotypic impact of such mutations is often milder than that of complete loss of function of the upstream transcription factor, kinase or phosphorylated target. Since the experimental design of the CGE assay mitigates the phenotypic effects associated to genome editing process itself, the assay is sensitive enough for detecting the subtle effects resulting from mutating transcription factor binding sites and posttranslational modifications. Here we identify several MYC binding motifs at the promoters of its target genes that are critical for cellular fitness. The critical target genes represent the major pathways previously associated to MYC function^6^: 1) ribosome biogenesis, including RPL23, a component of 60S large ribosomal subunit, and MDN1, a nuclear chaperone required for maturation and nuclear export of pre-60S ribosome subunit^37^, 2) cellular metabolism, as shown for glycolytic enzyme HK2, as well as 3) nucleotide synthesis, as shown for PPAT involved in *de novo* purine biosynthesis. Interestingly, however, mutation of the E-box at the SHMT2 promoter had no effect on cellular fitness in HAP1 cells, although SHMT2 has been previously shown to partially rescue the growth defects of Myc-null fibroblast cells^38^. These results highlight the importance of precise quantitative studies in determining the functional consequence of transcriptional regulatory events on cellular phenotype.

In summary, we report here an advanced method for measuring the phenotypic effects of precise targeted mutations. The method allows controlling for the effect of DNA damage, the major confounder in CRISPR-based methods. We also demonstrate the power of the technology by robustly detecting small fitness effects of individual transcription factor binding motifs and single amino acid substitutions. The method is widely applicable and extends the utility of CRISPR/Cas9-mediated genome editing to address important biological questions that have been difficult to address using existing technologies. Using this technology, we identified several target genes whose regulation via canonical E-boxes is responsible for the growth-promoting activity of the universal oncogene MYC.

## Methods

### Genome editing constructs

Precision editing of each genomic locus was performed by introducing a CRISPR/Cas9-mediated DSB and HDR templates harboring either the mutated or original genomic feature along with a sequence tag. Guide sequences were designed using CRISPOR^39^ tool, giving preference to the protospacers with closest distance to the genomic feature to be edited, and the crRNAs were obtained from Integrated DNA Technologies (**Supplementary Tables 1, 2**). Single-stranded 100-nucleotide (nt) DNA molecules were used as HDR templates. For editing E-box sequences, two HDR templates were designed for each targeted locus, one with CACGTG sequence for reconstituting the original E-box, and another with mutated sequence that replaces the E-box with non-functional TATTTA sequence. In each oligo, the original or mutated E-box was flanked by a 10-nt sequence tag and two 42-nt homology arms complementary to the target strand. Sequence tags were generated by mutating each of the ten nucleotides with probability of 24%, i.e. 8% probability for each of the three non-consensus bases (oligo synthesis using custom hand-mixed bases from Integrated DNA Technologies; **Supplementary Table 1**). As a negative control for E-box mutation experiments, the coding region of the *MYC* gene was targeted with two HDR templates, one reconstituting the original coding sequence, and another replacing nucleotides encoding Val-5 and Ser-6 with synonymous codons (GTTAGC > GTAAGT), and the sequence tag was created by randomizing the third degenerate position in the two codons flanking the targeted region on both sides. For targeting protein phosphorylation sites in the coding regions of the *CDK1* and *GRB2* genes, HDR oligos with 40-nt homology arms and sequence tags generated by randomizing the degenerate positions of the adjacent codons in the repair template were designed (**Supplementary Table 2**). Phosphorylation sites were abolished by alanine or phenylalanine substitutions of the phosphorylated serine/threonine or tyrosine residue, as well as mutated to glutamate or aspartate that in some cases mimic the phosphorylated state^22^.

Prime editing guides (pegRNA) to target CDK1 were designed according to the recommendations from ref. ^30^. Similar to HDR templates described above, the pegRNA pool introduces a mutation (Y15F) or reconstitutes the original sequence, and in both cases the third degenerate position in the codons flanking the targeted region was randomized to create the sequence tags for lineage-tracing (**Supplementary Table 2**).

PCR primers for amplifying genomic DNA at each targeted locus were designed so that neither forward nor reverse primers overlap with the genomic sequence used in the HDR templates. All custom oligoes used for targeting and amplifying the E-box targets and the phosphorylation sites are listed in **Supplementary Tables 1** and **2**, respectively.

### Cell lines and transfections

HAP1 (#C631) and KMB-7 (#C628) cell lines were obtained from Horizon Discovery and maintained in low-density cultures in Iscove’s Modified Dulbecco’s Medium (IMDM) with 10% FBS, 2 nM L-glutamine, and 1% antibiotics according to the vendor’s guidelines.

Precision editing experiments measuring cellular fitness were done by transfecting 200,000-400,000 early-passage HAP1 or KBM-7 cells with ribonucleoprotein (RNP) complex and the two HDR templates. For sgRNA molecules, equimolar ratios of target-specific crRNAs and ATTO550-tracrRNA (Integrated DNA Technologies) were annealed. RNP complexes used for the transfections were constituted from S.p. HiFi Cas9-protein (Integrated DNA Technologies; 1000 ng / 200,000 cells) and target-specific sgRNA (250 ng / 200,000 cells) and transfected to cells using CRISPRMAX (Life Technologies) as per manufacturer’s recommendation along with HDR template (1:1 mixture of the original and mutant HDR templates) with final concentration of 3 nM. Half of the cell population was harvested for genomic DNA isolation 48 h after transfection (day 2), and the rest of the cells were plated for culture and harvested on day 8. For ChIP assays measuring the effect of E-box mutation on MYC occupancy and H3K27 acetylation, 15 million cells were transfected for each condition on two 15-cm dishes, scaling up the transfection reagents according to the cell numbers. The cells were harvested, and chromatin cross-linked 48 h after transfection.

For prime editing experiments, Prime editor 2 was expressed from pCMV-PE2 and pegRNAs from pU6-pegRNA-GG-acceptor plasmids^30^ (Addgene #132775 and #132777, respectively). Plasmid transfection was performed using FuGENE HD (Promega) according to manufacturer’s instructions. Rest of the experiment was performed in the same way as the homology-directed repair editing experiment.

### Genomic DNA isolation and target-specific sequencing

Genomic DNA was isolated using AllPrep DNA/RNA Mini kit and Blood & Cell Culture DNA Maxi kit Qiagen) from day 2 and day 8 time points, respectively, and treated with RNAse A (0.2 μg/ul; Thermo Fisher Scientific) for 2 h at 37 C. To eliminate the carry-over of single-stranded DNA from the HDR templates, gDNA samples were treated with exonuclease I and VII (New England Biolabs) in 10 mM Tris-HCl, 50 mM KCl, 1.5 mM MgCl2 for 30 min at 37 C followed by enzyme inactivation for 10 min at 95 C and DNA extraction using phenol:chloroform:isoamyl alcohol (Sigma). All gDNA from day 2 and 10 μg gDNA corresponding to 3 million haploid cells from day 8 was amplified in PCR reactions with maximum of 2.5 μg of DNA per reaction using NEBNext High Fidelity master mix (New England Biolabs) and target-specific primers with Illumina adaptor flanks (**Supplementary Tables 1, 2**) for 20 cycles, followed by DNA purification using 1.5x Ampure XP beads (Beckman Coulter). For amplifying the genomic regions with E-box targets, biotinylated primers were used for PCR1, and the 30% volume of the purified PCR products was used for streptavidin capture with M-280 Dynabeads (Thermo Fisher Scientific) according to manufacturer’s protocol. Prime-editing samples are not affected by the presence of HDR template, and thus they were prepared without the exonuclease I/VII treatment and affinity purification of biotinylated PCR products. Second PCR amplification with 8 cycles was used for generating sequencing-ready libraries using NEBNext High Fidelity master mix and Illumina Universal and Index primers (E7335S, New England Biolabs) in four and twelve parallel reactions on M-280 beads for day 2 and day 8 samples, respectively. PCR products were purified using 0.9x Ampure XP beads and sequenced for 150 cycles on NovaSeq 6000, HiSeq 4000, and NextSeq 500 platforms (Illumina).

### Analysis of precision editing data

Raw sequencing files were demultiplexed with bcl2fastq software. Sequencing reads were assigned to each genomic target by matching the first 20 nucleotides of each read to the sequence of PCR products amplified using target-specific primers. Then, the reads originating from cells that had undergone successful precision editing were identified based on the presence of a sequence tag, and the reads originating from un-edited wild-type cells were discarded. Reads matching to each individual sequence tag flanking the original and mutated sequence features were counted in different timepoint samples, and a pseudocount of +1 was added to each read count to avoid the zeros in subsequent calculations. For experiments targeting E-boxes, all cell lineages harboring sequence tags with read count > 50 in both day 2 samples were included in the analyses. In **Fig. 1d**, the results from all sequence tags with exactly one flanking mutation are shown to demonstrate the power of the assay in tracing the growth of individual cell lineages over time. However, to further increase the robustness of the analysis, only the sequence tags with exactly two flanking mutations were included in the analyses for **Fig. 2b, c** and **Supplementary Fig. 4**. For experiments targeting protein phosphorylation sites, all the sequence tags with read count > 20 for GRB2 experiments and > 5 for CDK1 experiments were included in the analyses due to lower sequencing depth in these experiments.

To analyze the effect of each mutation on cellular fitness, the ratio of cells harboring mutated and original sequence features was compared at each time point. If mutating a particular E-box or phosphorylation site hampers cell growth or proliferation, the cells harboring the mutated allele will be underrepresented in the final pool of cells after one-week culture compared to the cells harboring the original sequence feature, and vice versa. Thus, the read count ratios were calculated for each sequence tag for mutated vs. original sequence. To eliminate the potential effect of near-silent flanking mutations on cellular fitness, the sequence tags with the same flanking mutations were compared in the analysis. Finally, the ratio for day 8 vs. day 2 was calculated for each sequence tag. In **Fig. 2**, the results are presented as log_2_ (fold change) as follows: log_2_[day 8 read count (mutated / original) / day 2 read count (mutated / original)] for each cell lineage. Wilcoxon signed rank test was used for testing whether the median of log_2_ (fold change) values is unequal to zero.

### Chromatin-immunoprecipitation (ChIP) with target-specific sequencing and ChIP-seq

Wild-type HAP1 cells and genome-edited HAP1 cells 48 h after RNP transfection were crosslinked with 1% formaldehyde and chromatin samples were prepared as described previously^40^. Chromatin was sonicated to an average fragment size of 500 bp using micro-tip sonicator (Misonix Inc) and used for immunoprecipitation (IP) with antibody-coupled Dynal-beads (Thermo Fisher Scientific) for MYC, H3K27ac, and normal rabbit IgG (Millipore #06-340, Abcam #ab4729, and SantaCruz #sc-2027, respectively). Chromatin from ten million wild-type cells and 20 million transfected cells was used for each IP. After overnight incubation, washed with LiCl buffer and reverse cross-linking was performed as described in ref. ^40^, followed by DNA purification using phenol:chloroform:isoamyl alcohol and ethanol precipitation.

All immunoprecipitated DNA isolated from transfected cells was amplified for 30 cycles in two reactions using similar PCR strategy and conditions as described above for genomic DNA samples. In addition, 10 μg of input DNA from each transfected condition was similarly amplified in four parallel reactions. PCR products were purified using 1.5x Ampure XP beads and 20% of purified DNA was used as a template for the second PCR amplification step for 8 cycles with Illumina primers as above. Final libraries were purified using 0.9x Ampure XP beads and sequenced for 150 cycles on NovaSeq 6000 (Illumina). Data was analyzed essentially as described for fitness experiments: after excluding the reads originating from wild-type cells, a pseudocount +1 was added to the reads originating from the edited cells with distinct sequence tags, the read count ratios between mutated and original sequences were calculated for each condition, and log_2_(fold change) between each IP and respective input sample was calculated. Only cell lineages harboring sequence tags with read counts > 50 in the input sample were included in the analyses. Wilcoxon signed rank test was used for testing whether the median of log_2_ (fold change) values is unequal to zero.

Wild-type HAP1 samples were used for standard ChIP-seq library preparation with NEBNext Ultra II DNA Library Prep kit (New England Biolabs), followed by sequencing on NovaSeq 6000. The reads were aligned to human genome (hg19) using bowtie2^41^ and peaks were called using MACS2^42^ against input with default narrow peak parameters. The bedgraph files were used for genome browser snapshots. For colon cancer cell lines GP5d, LoVo, and COLO320DM, previously published ChIP-nexus data sets from ref. ^32^ (EGAD00001004099) were used. In the genome browser snapshots, the traces from BAM coverage files are shown.

### Chromatin accessibility and gene expression analysis

ATAC-seq for chromatin accessibility was performed from 50,000 HAP1 cells as previously described^43^. Briefly, the cells were washed with ice-cold PBS, lysed in 50 μl of lysis buffer for 10 min in ice, and treated with Tn5 transposase in 2x tagmentation buffer (Illumina) for 30 min at 37 C. DNA was purified using MinElute PCR Purification kit (Qiagen) and prepared for sequencing using Nextera library preparation kit (Illumina) by five cycles of PCR amplification. The sample was sequenced on NovaSeq 6000 for 2 x 50 cycles, and the paired-end data was analyzed using an in-house pipeline as described in ref. ^33^. For GP5d cells, the ATAC-seq data from ref. ^33^ (GSE180158) was used. In the genome browser snapshots, the traces from BAM coverage files are shown.

For gene expression analysis, previously published RNA-seq data from ref. ^32^ (EGAD00001004098) was used. The data sets for MYC silencing using siRNA (siMYC) and respective control samples transfected with non-targeting siRNAs (siNon-target) for GP5d and LoVo cells were re-analyzed by aligning the reads from fastq files to human genome (hg19) using tophat2^44^ and by analyzing the differentially expressed genes between siMYC and siNon-target samples using cuffdiff^45^ using default parameters and the option for first-strand library type. Log_2_(fold change) values for the selected genes are shown in **Supplementary Fig. 3**.

## Acknowledgements

We thank Anu M. Luoto, Kaisu Jussila, and Katariina Sarin for technical assistance. We also thank HiLIFE research infrastructures including the Biomedicum Functional Genomics Unit (FuGU) and the Sequencing laboratory of Institute for Molecular Medicine Finland FIMM Technology Centre, University of Helsinki. JT was supported by grants from the Academy of Finland (Finnish Center of Excellence program: 2012-2017, 250345 and 2018-2025, 312042), Finnish Cancer Foundation, and Cancer Research UK (RG99643). PP was supported by the Academy of Finland (288836). BS was supported by the Academy of Finland (274555, 317807), Finnish Cancer Foundation, Sigrid Jusélius and Jane and Aatos Erkko Foundations.

## Author contributions

JT conceived and supervised the study. PP designed and performed E-box editing experiments and analyzed the data with conceptual help from TK. OK designed, performed and analyzed editing experiments for the phosphorylation sites with input from PP. BS performed and analyzed the ChIP-seq and ATAC-seq experiments. All authors contributed to the writing of the manuscript.

## Corresponding author

Correspondence and requests for materials should be addressed to

## Data availability

All next-generation sequencing data generated in this study is available under ENA accession XXX. Previously published data sets for colon cancer cells were used as follows: RNA-seq from EGAD00001004098, ATAC-seq from GSE180158, and ChIP-nexus from EGAD00001004099.

